# Biochemistry-informed design selects potent siRNAs against SARS-CoV-2

**DOI:** 10.1101/2022.12.08.519651

**Authors:** Élisabeth Houbron, Sophie Mockly, Sophia Rafasse, Nathalie Gros, Delphine Muriaux, Hervé Seitz

## Abstract

RNA interference (RNAi) offers an efficient way to repress genes of interest, and it is widely used in research settings. Clinical applications emerged more recently, with 5 approved siRNAs (the RNA guides of the RNAi effector complex) against human diseases. The development of siRNAs against the SARS-CoV-2 virus could therefore provide the basis of novel COVID-19 treatments, while being easily adaptable to future variants or to other, unrelated viruses. Because the biochemistry of RNAi is very precisely described, it is now possible to design siRNAs with high predicted activity and specificity using only computational tools. While previous siRNA design algorithms tended to rely on simplistic strategies (raising fully complementary siRNAs against targets of interest), our approach uses the most up-to-date mechanistic description of RNAi to allow mismatches at tolerable positions and to force them at beneficial positions, while optimizing siRNA duplex asymmetry. Our pipeline proposes 8 siRNAs against SARS-CoV-2, and *ex vivo* assessment confirms the high antiviral activity of 6 out of 8 siRNAs, also achieving excellent variant coverage (with several 3-siRNA combinations recognizing each correctly-sequenced variant as of September 2022). Our approach is easily generalizable to other viruses as long as a variant genome database is available. With siRNA delivery procedures being currently improved, RNAi could therefore become an efficient and versatile antiviral therapeutic strategy.

## INTRODUCTION

RNA interference (“RNAi”) is an efficient repressive mechanism broadly conserved among eukaryotes. While there exists a diversity of clade-specific features [1], the core mechanism of RNAi always involves small RNA guides (“siRNAs”) loaded on a protein of the AGO subfamily of the Argonaute family [2]. The resulting ribonucleoprotein complex (the “RISC” complex) then binds target RNAs exhibiting sufficient sequence complementarity, and cleaves them endonucleolytically between the nucleotides facing guide nucleotides 10 and 11 [3, 4, 5, 6, 7]. Mismatches in the vicinity of the scissile phosphate tend to inhibit the cleavage reaction [4, 8, 9, 10, 11].

siRNAs are double-stranded (with each strand being ≈21 nt long). After loading on the AGO protein, one strand (the “passenger strand”) is discarded while the other one (the “guide strand”) remains stably associated with AGO. Mature RISC therefore contains a single-stranded small RNA, which is available for base-pairing with target RNAs [12]. Among the four existing AGO’s in mammals, only one (named “AGO2”) possesses a competent catalytic site for robust cleavage activity [13, 14, 15, 16].

Besides RNAi, other related pathways also implicate small RNA-loaded Argonaute proteins. In particular, the microRNA (“miRNA”) pathway, where the small guide RNA follows a distinctive biogenesis route before being loaded on an AGO protein [1]. In bilaterian animals, miRNAs typically recognize their targets by imperfect sequence complementarity: in general, the target exhibits a perfect match to the miRNA “seed” (nt 2–7), but mismatches are well tolerated at most other positions [17]. Just like siRNAs, miRNAs can guide target cleavage if sequence complementarity requirements are met [18], but this is very rare in bilaterian animals [19, 20, 21, 22]. If conditions are not met for target cleavage (either because of mismatches close to the scissile phosphate, or because the AGO protein does not possess a catalytic site for cleavage), the target RNA is still repressed, but less efficiently, by various mechanisms (including exonucleolytic degradation and, for mRNAs: translational inhibition; [23]).

The efficiency and specificity of RNAi make it a powerful tool for targeted gene repression in laboratory settings. It has also quickly been proposed to serve as a basis for antiviral therapy [24] but clinical applications have been slower, with only 5 approved therapeutic siRNAs so far, none of them targeting a virus [25, 26, 27, 28, 29]. Obstacles to the development of therapeutic siRNAs include the poor stability of RNA in serum, and the tendency for injected siRNAs to concentrate primarily in the liver. Efforts for the development of these five approved drugs resulted in the discovery of chemical modifications which greatly stabilize siRNAs *in vivo* while not compromising too much their activity. Regarding pulmonary diseases, the biodistribution issue may also be circumvented by the development of inhalation-based procedures [30].

The coronavirus disease 2019 (COVID-19) outbreak started during the winter 2019–2020 and it was initially detected in Wuhan (China) [31]. The causative viral agent, the severe acute respiratory syndrome coronavirus 2 (SARS-CoV-2), was quickly identified and its genome was sequenced from patient samples and distributed via the GISAID repository [32]. Hundreds of viral variants from individual donors were available in the first few weeks of the pandemic (see Figure 1**A**). Because the biochemistry of RNAi is well known, it is possible to design siRNA sequences with predicted high efficacy — and because so many sequenced viral variants were quickly available, it is possible to choose siRNA sequence sets which should be active against every known variant. Here we describe our bioinformatics approach for the design of siRNAs against SARS-CoV-2, and verification of their activity in a simple, SARS-CoV-2 infected cultured cell-based model *ex vivo*. A comparison with previously published works shows that our purely computational design yields a high proportion of efficient siRNAs against SARS-CoV-2 that were validated *ex vivo*. Once delivery methods have been optimized for *in vivo* administration, antiviral siRNAs could therefore offer a quick response, easily designed and easily reprogrammable against novel variants, or other viruses in the case of future viral pandemics.

**Figure 1.**
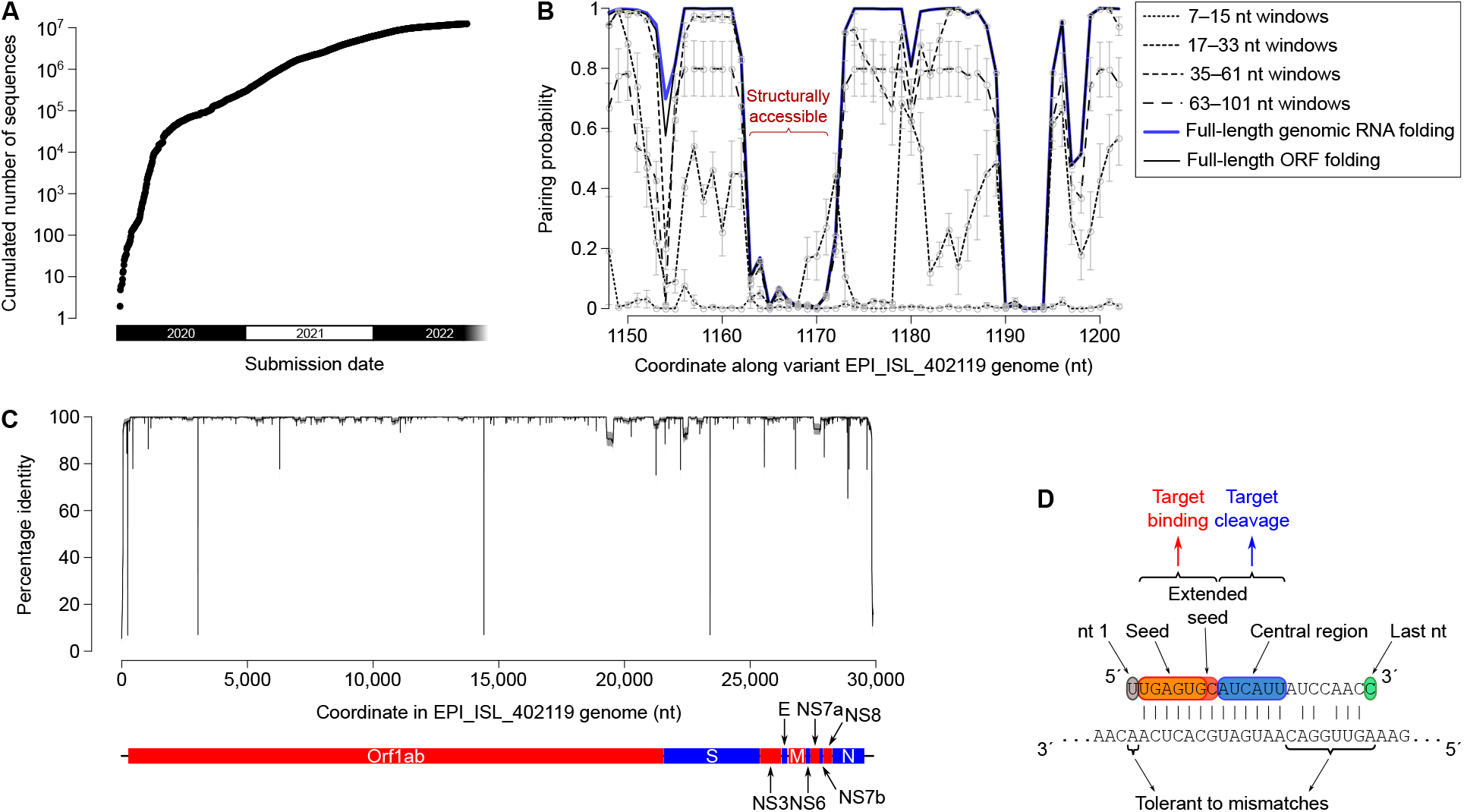
Criteria for siRNA design. **A**. Cumulated number of SARS-CoV-2 variant sequences from human donors at GISAID as a function of time. **B**. Predicted structural accessibility of nt 1150–1200 in viral variant #EPI ISL 402119 (a.k.a. BetaCoV/Wuhan/IVDC-HB-01/2019). *x* axis: genomic coordinate. *y* axis: nucleotide pairing probability in various sequence contexts (full-length genomic RNA folding, full-length ORF folding, and various sliding window size ranges). Gray circles represent the mean pairing probability across window sizes and gray error bars represent the standard error. The red curly bracket indicates a structurally accessible region. **C**. Nucleotide conservation along EPI ISL 402119 genome sequence. 100 non-overlapping sets of 100 variants each were randomly picked among the 311,994 correctly-sequenced viral variants from human donors in GISAID release dated Jan. 8, 2021, then each of these sets was aligned with the EPI ISL 402119 sequence. Identity to EPI ISL 402119 was scored for each nucleotide in that reference sequence; the black curve represents the mean, and the gray shading represents the mean +/-standard deviation, across all 100 random sets. Annotated ORFs in the EPI ISL 402119 genome are represented under the graph (red and blue bars). **D**. Functional modules in a siRNA guide strand sequence (top strand) for the recognition and cleavage of a target RNA (bottom strand). Guide nt 2–7 constitute the “seed” and nt 2–8 constitute the “extended seed” [17, 49]. Guide nt 9–13 (and 9–14 for some siRNA sequences) need to be perfectly paired to the target for optimal cleavage rate according to Cleave’n-Seq experiments [7]. Other positions are more tolerant to mismatches without compromising target binding or cleavage. The last nucleotide is preferentially unpaired (respectively to its target) in order to accelerate product release and reduce the risk of TDMD-mediated siRNA destabilization or guide strand unloading [11, 50, 7].

## MATERIALS AND METHODS

### Computational analyses

Viral variant sequences were downloaded from the GISAID database (https://gisaid.org/). Conserved regions were identified from **clustalw**-generated alignments of correctly sequenced variant sequences (at least 26 kb long, no more than 5% “N” ambiguities). RNA secondary structures in viral RNAs were predicted using **RNAfold** on a reference variant sequence (#EPI ISL 402119).

Scripts, raw, intermediate and final data files are available at https://github.com/HKeyHKey/Houbron_et_al_2023 and at https://www.igh.cnrs.fr/en/research/departments/genetics-development/systemic-impact-of-small-regulatory-rnas#programmes-informatiques/.

### Cell culture, transfection and assessment of siRNA cytotoxicity

Vero E6 cells (ECACC batch #15A033) were grown in DMEM (with 10% FBS, 1% L-glutamine, 1% penicillin+streptomycin, no glutamax; Gibco) at 37^*?*^C under 5% CO_2_. siRNAs were transfected using Lipofectamine RNAimax (Invitrogen) according to the manufacturer’s protocol, 24 h after seeding 96 well-plates with 3.75 ×10^3^ cells per well (for the siRNA cytotoxicity assay) or 3× 10^4^ cells per well (for antiviral assessment by RT-qPCR or cytopathic effect measurement). siRNA cytotoxicity was assessed 72 h post-transfection using the CellTiter Glo kit (Promega). For A549-hACE2 cells (strain described in [33, 34]), protocols were identical except that the cells were grown in RPMI with L-glutamine (with 10% FBS, 1% penicillin+streptomycin; Corning), transfected with Lipofectamine 3000 (Invitrogen) according to the manufacturer’s protocol, and seeded at 2.5 ×10^3^ cells per well for the siRNA cytotoxicity assay, and 2 ×10^4^ cells per well for the antiviral activity assay.

### Viral infection and assessement of antiviral activity

Six hours after siRNA transfection, the medium was removed, cells were rinsed with PBS, then incubated in fresh DMEM (for Vero E6) or RPMI (for A549-hACE2) medium. One hour later, SARS-CoV-2 (strain #2020-A0093534 isolated by CPP Île de France III and provided by Collection resource CRB of the Montpellier hospital) was inoculated at MOI=0.01. Cytopathic effect was assessed using the Viral ToxGlo Assay kit (Promega) 72 h after infection. For viral RNA quantification: cells were rinsed with PBS, lysed with the Luna Cell Ready Lysis Module (New England BioLabs): in a 96 well plate, cells were lysed in 40 *µ*L lysis buffer per well (10 min incubation at 37^*?*^C, then 5 *µ*L Stop mix was added to each well), then lysates were transferred to a new plate and frozen at -80^*?*^C. 1 *µ*L lysate was used as a template for the Luna Universal One-Step RT-qPCR Kit (New England Biolabs) on a Light Cycler 480 (Roche). Viral RNA (mRNA for gene E) was amplified with primers ACAGGTACGTTAATAGT-TAATAGCGT and ATATTGCAGCAGTACGCACACA; for normalization, cellular GAPDH mRNA was amplified with primers GCTCACTGGCATGGCCTTCCGTG and TGGAGGAGTGGGTGTCGCTGTTG.

As a positive antiviral control, Remdesivir was added to the cell medium at 10 *µ*M after viral infection.

### Ethics

Ethical approval was obtained from the cell culture room’s institutional review board and ethics committee (protocol number 2010-A01406-33) as is classically done in such studies.

## RESULTS

### Rationale for siRNA design

Biochemical determinants of siRNA activity are now very well described, therefore theoretically offering the possibility to design efficient siRNAs with a pure computational approach. Because SARS-CoV-2 is a positive-strand RNA virus, siRNAs raised against genomic RNA can also target viral mRNAs, while siRNAs against the negative strand would only target short-lived replication intermediates, resulting in a lower efficiency [35]. We therefore applied the following criteria for the design of positive strand-targeting siRNAs against SARS-CoV-2:

- Target sites are poorly accessible to siRNAs if they can fold on themselves, forming stable intramolecular stems [36]. Target secondary structure can be predicted by conceptual folding of the full-length genomic RNA, full-length mRNAs (here approximated by their ORF’s), or by shorter sliding windows (short-range interactions are expected to re-form rapidly after ribosome scanning). We therefore selected siRNA candidates targeting viral regions which are predicted to be accessible according to each of these metrics (see Figure 1**B**).
- Viral genomes tend to mutate very quickly, and it can be anticipated that, in the long-term, an siRNA treatment would promote the appearance of resistant variants (where the siRNA target site has been mutated). We therefore selected siRNA candidates which can target the most conserved viral regions: not only they should be active against already-existing variants, but likely also against future variants (strong sequence conservation in the viral genome marks functionally important segments, which could not mutate without compromising viral spread) (see Figure 1**C**).
- If imperfect base-pairing between the siRNA guide strand and the target cannot be avoided (which is the case when designing siRNAs against a large diversity of viral variants), mismatches may be harmlessly introduced at positions 1, 15, 17, 18, 19, 20 or 21 of the guide RNA [37, 38, 7]. Mismatches at other positions often reduce binding affinity of RISC for its target, or the catalytic rate of the cleavage reaction — sometimes in a sequence-dependent manner. The first nucleotide of the siRNA guide strand does not need to be complementary to the facing target nucleotide: it is anyway unpaired in the RISC complex, even when the facing nucleotide is complementary [39]. We therefore selected 13-mer target regions (to be targeted by siRNA guide nt 2–14) which are conserved among viral variants (see Figure 1**D**).
- Human AGO2 binds preferentially small RNAs with a 5’ uridine or a 5’ adenosine [40]. And more precisely, natural human miRNAs frequently have a 5’ uridine [41], which may be due to an intrinsically higher affinity of the AGO protein or its loading machinery. We therefore introduced a 5’ uridine in each siRNA candidate.
- In addition to the intended target, introduced siRNAs are likely to bind additional mRNAs (“off-targets”): these are human endogenous mRNAs with fortuitous binding sites for the siRNA. If there are many off-targets, the siRNA is likely to be partially titrated, hence less efficient [42]. And because off-targets are (moderately) repressed by the siRNA, they could trigger unwanted secondary effects [43]. The most efficient binding sites (“7mers” and “8mers”; [17]) are so short that it is impossible to avoid off-targets (some human mRNAs will unavoidably contain perfect seed matches in their 3’ UTR). But it is possible to minimize the number of off-targets, and to minimize the number of off-targets whose modest down-regulation is most susceptible to trigger phenotypic consequences in humans (here approximated by a curated list of human haplo-insufficient genes; [44]).
- In an siRNA duplex, differential base-pairing stability on the two ends of the duplex principally determines the identity of the guide strand [45, 46, 47]. Fraying the 5’-most nucleotide of the intended guide strand, and favoring candidate siRNAs with intrinsic asymmetry (high AU-content at the 5’ end of the intended guide strand, high GC-content at the 5’ end of the intended passenger strand) therefore improves siRNA efficiency.
- Once the RISC complex is matured, it is more stable if the guide strand contains a 5’ phosphate [15]. Functional asymmetry of the duplex can therefore be further improved by phophorylating the 5’ end of the intended guide strand, while the 5’ end of the intended passenger strand is unphosphorylated [48].

Our siRNA design workflow is presented in Figure 2**A**. It resulted in the selection of 8 target 13-mers whose seed match is structurally accessible according to every metrics, which are perfectly conserved in *>*99.8% correctly-sequenced viral variants, and which tend to minimize the number of haplo-insufficient predicted off-targets (see Figure 2**B**). Even though our method was applied to the whole viral genome, each of these 8 candidate target sites is located in the Orf1ab coding sequence of the viral genome (see below, section “Comparison to previously-published siRNAs and recent viral variants”). This is probably due to the fact that Orf1ab is the longest gene in SARS-CoV-2 (accounting for more than two-thirds of the total genome length), as well as to the presence of long stretches of highly constant regions coding for viral enzymes expressed from that gene. siRNA duplexes against these 8 target sites are presented in Figure 2**C**: they were optimized for their functional asymmetry (with the intended guide strand being 5’-phosphorylated, and having less stably paired 5’ nucleotides than the intended passenger strand), and the 3’-most nucleotide of their intended guide strand cannot pair to the facing nucleotide in the viral RNA target. Fraying the guide’s 3’ end when paired to its target indeed accelerates product release and reduces the risk of guide strand destabilization by TDMD (target-directed microRNA degradation) or by AGO unloading [11, 50, 7].

**Figure 2.**
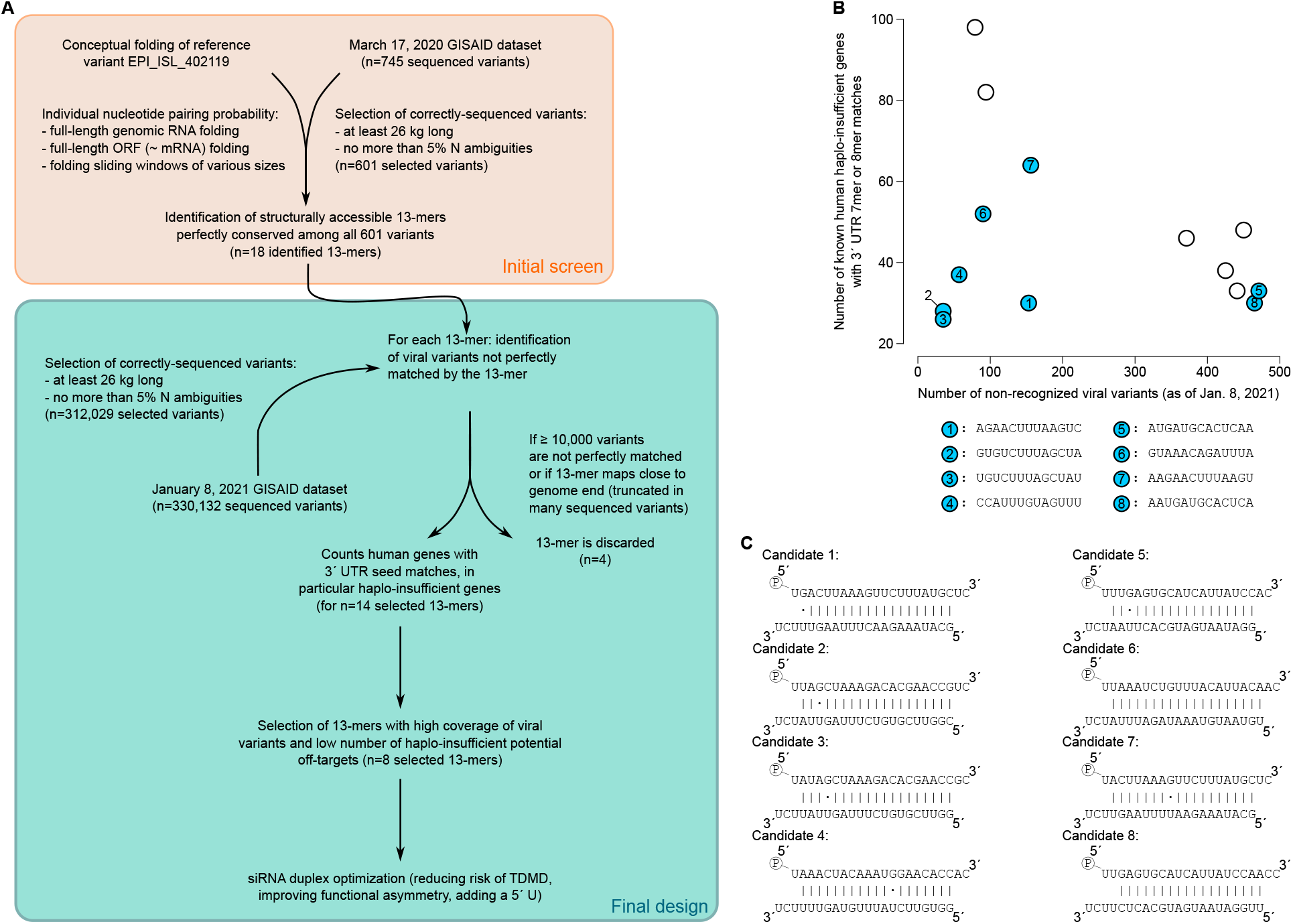
siRNA selection. **A**. Workflow for siRNA design. An initial screen (orange frame) identified a list of 18 candidate 13-mer structurally-accessible target sites conserved among sequenced variants as of March 17, 2020. Final siRNA design screened these 18 candidates for their conservation in an updated variant list (as of January 8, 2021) and ranked them by their number of un-recognized viral variants and their number of predicted haplo-insufficient off-targets. **B**. Final choice of 8 target 13-mers. *x*-axis: number of non-recognized viral variants (*i*.*e*.: viral variants without a perfect 13-mer match not due to sequence ambiguity) out of 312,029 variants in total. *y*-axis: number of haplo-insufficient predicted human off-targets (*i*.*e*.: bearing a 7mer or 8mer site for the siRNA seed in their 3’ UTR). Eight 13-mers with low numbers of missed viral variants and low numbers of predicted off-targets were selected and siRNAs were raised against each (these 13-mers are colored in blue, their sequences are given under the graph). **C**. Optimized siRNA duplexes targeting the 8 selected 13-mer targets. In each duplex, the intended guide strand is at the top and the intended passenger strand is at the bottom.

### *Ex vivo* assessment against SARS-CoV-2

The eight candidate siRNAs were synthesized and transfected in Vero E6 cells for cytotoxicity assessment. Vero E6 is a fetal *Cercopithecus aethiops* monkey kidney cell line particularly permissive to replication of SARS-CoV-2, and where the virus causes a strong cytopathic effect [51]. In the conditions tested, none of the siRNAs triggered significant cell death 72 h after transfection. Three candidates (siRNAs #1, 5 and 6) actually increased cell proliferation at high concentration (100 nM), with one of them (siRNA #6) also significantly increasing proliferation at 20 nM (see Figure 3**A**), which remains unexplained. We also verified the absence of siRNA-induced lethality in a human cell line (the lung epithelium-derived A549 line expressing a stable hACE2 transgene to make it susceptible to SARS-CoV-2 infection; [52]). In this human-derived cell line too, none of the 8 siRNAs induced cell death (see Supplementary Figure 1**A**).

Antiviral activity was then evaluated by transfecting 20 nM siRNAs in Vero E6 cells, then infecting the cells with SARS-CoV-2 seven hours post-transfection. The virus is not washed out letting the infection to propagate for 3 days. Viral RNA contained in the cells was quantified by RT-qPCR on gene E 48 h after infection, and the cytopathic effect (CPE) was quantified by measuring cell viability 72 h after infection. These two independent measurements gave similar conclusions, with siRNAs #1 and 7 showing no clear antiviral activity, siRNA #6 showing a moderate activity, while siRNAs #2, 3, 4, 5 and 8 showed both a strong reduction in viral RNA titer and in CPE (see Figure 3**B** and **C**). Cytopathic effect was also assessed after transfection of various siRNA concentrations, followed by viral infection: results are very similar to the 20 nM siRNA condition (see Supplementary Figure 2). In the A549-hACE2 cell line too, the same 6 siRNAs exhibited high anti-viral activity as judged by RT-qPCR on gene E (see Supplementary Figure 1**B**).

**Figure 3.**
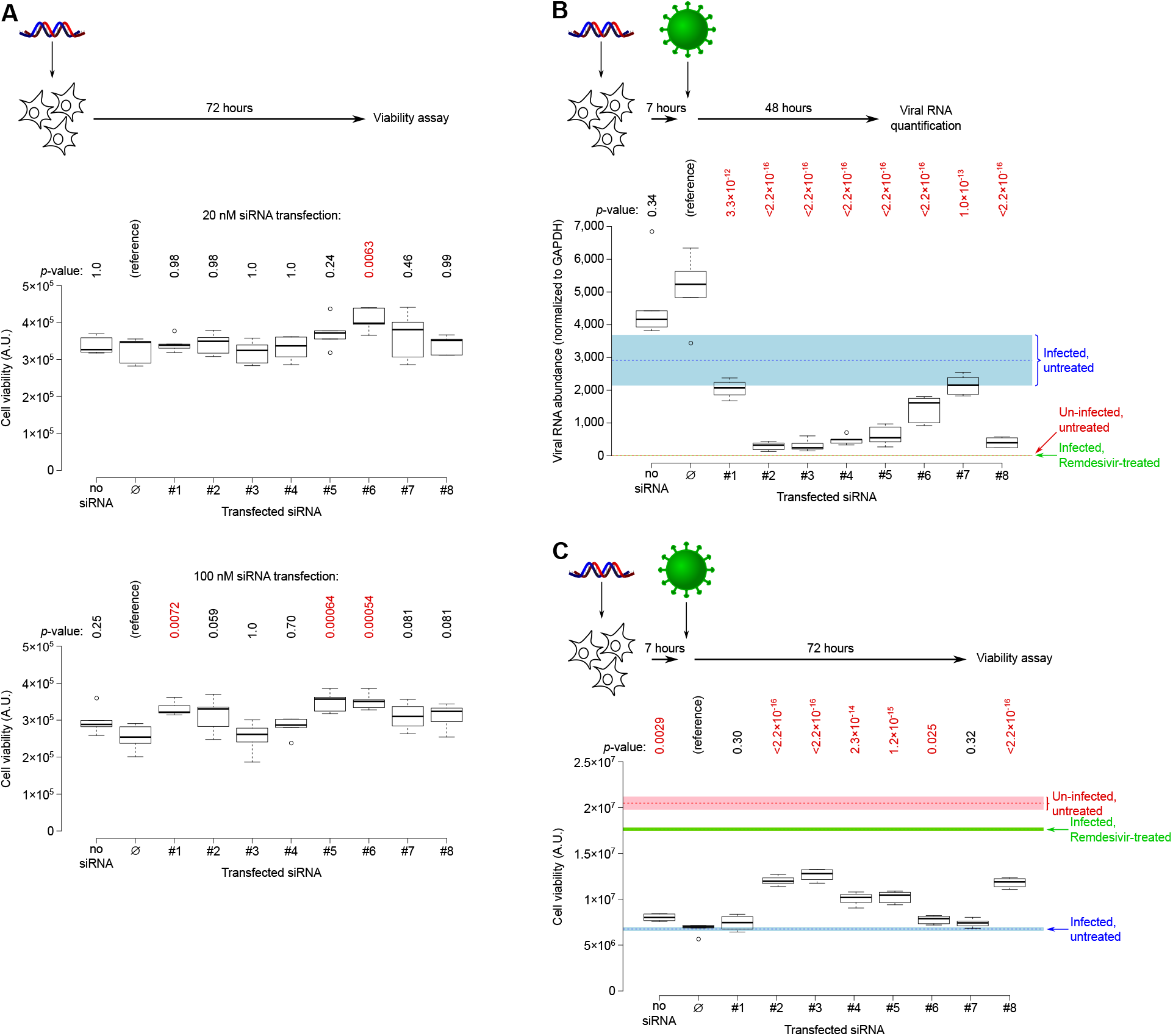
*Ex vivo* assessment of siRNA cytotoxicity and antiviral activity. **A**. Cell viability was measured 72 h after siRNA transfection (5 biological replicates). **B and C:** Cells were transfected with 20 nM siRNA and infected 7 h later (2 replicates for “Un-infected, untreated” and for “Infected, Remdesivir-treated”, an antiviral positive control; 4 replicates for “Infected, untreated”; 6 replicates for other conditions). Colored rectangles indicate the mean +/-standard error across replicates for infection and treatment controls. **B**. Viral RNA was quantified by RT-qPCR 48 h after infection, using endogenous GAPDH mRNA for normalization. **C**. Cell viability was measured 72 h after infection (see Supplementary Figure 2 for additional siRNA concentrations). **A, B and C:** Controls: “no siRNA” were transfected with empty Lipofectamine; “∅” were transfected with a control siRNA (designed to minimize its number of seed matches in mammalian genomes, while not having any 13-mer match to any sequenced SARS-CoV-2 variant as of Jan. 8, 2021). Significance was assessed using Dunnett’s test, with the “∅” siRNA as a reference; *p*-values are indicated above each graph, with *p*-values lower than 0.05 being shown in red.

A combination of two siRNAs (siRNAs #3 and 8) or of three siRNAs (siRNAs #3, 4 and 8) did not trigger significant cell death when transfected at a total concentration of 20 or 100 nM, either in Vero E6 or in A549-hACE2 cells (see Supplementary Figure 3**A** and **B**). Their anti-viral activity in Vero E6 cells was moderate (≈40% reduction in viral RNA) while it was similar to that of individual siRNAs in A549-hACE2 cells (92–95% reduction): see Supplementary Figure 3**C** and **D**.

### Comparison to previously-published siRNAs and recent viral variants

Several studies have already described the design and experimental evaluation of siRNAs against SARS-CoV-2, with some siRNAs being more active than others [53, 54, 55, 56, 57, 58, 59, 60, 61, 62, 35, 63]. Comparing success rates among diverse studies is not straightforward because of methodological differences between studies. To make the comparison as fair and uniform as possible, we compiled the results of *ex vivo* assessment of siRNAs against SARS-CoV-2 and flagged “high activity” those siRNAs with a significant repressive effect on viral RNA accumulation or viral plaque forming units exceeding 50% in the tested conditions (selecting conditions with siRNA concentrations close to 10 nM and post-transfection incubation close to 48 h whenever possible). The proportion of designed siRNAs with high activity ranges from 0 [57, 59] to 10/11 [61] and 1/1 [55] (see Figure 4; note that siRNAs designed by [58] could not be analyzed because they were tested *ex vivo* against SARS-CoV-2 only at a 100 nM concentration). Our approach, yielding 6/8 siRNAs with high activity, proved efficient in designing potent siRNAs with a pure bioinformatics approach, without requiring high-throughput experimental screening.

**Figure 4.**
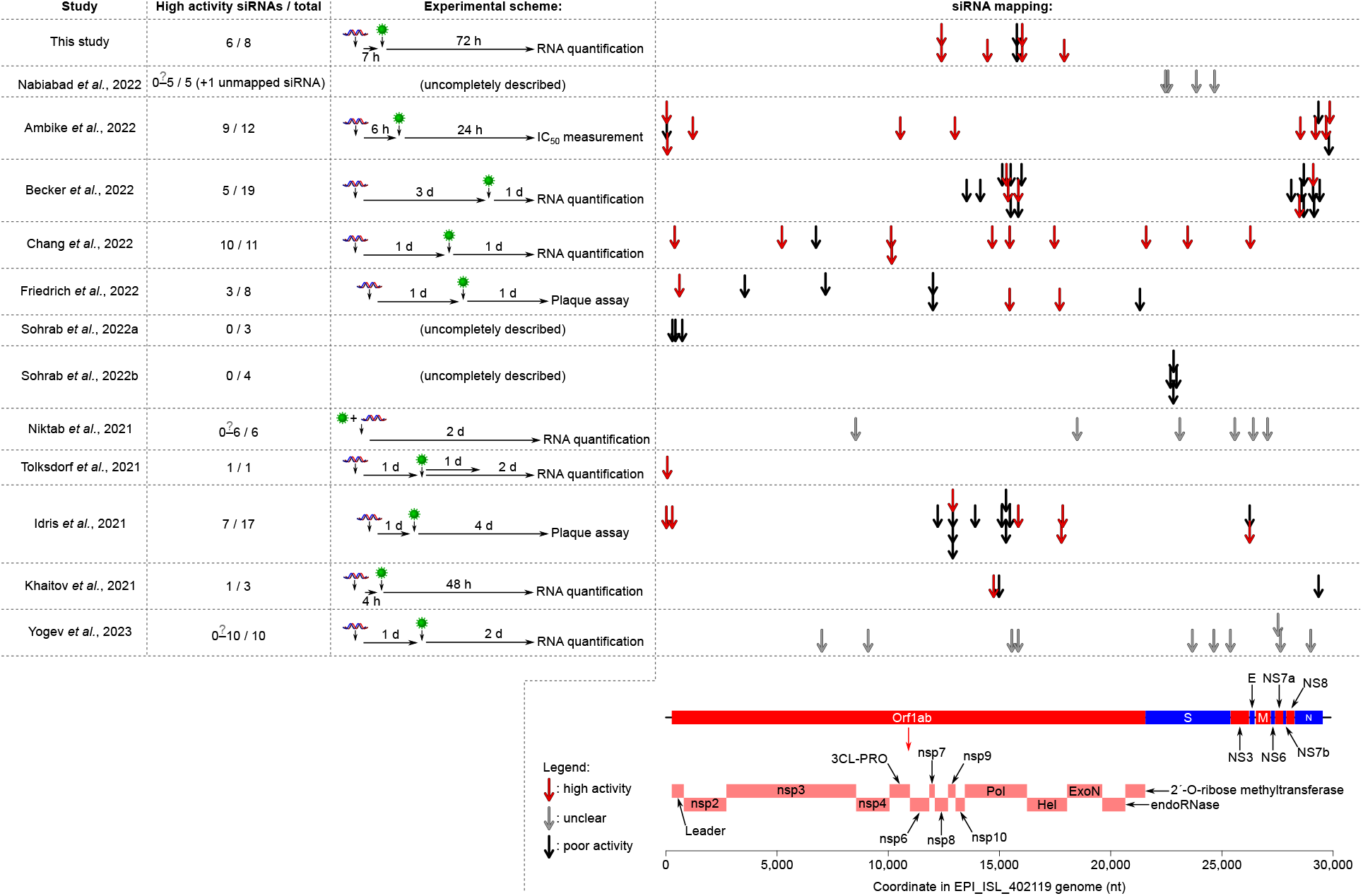
Comparison with previous anti-SARS-CoV-2 siRNA design attempts. Each arrow represents the target site for a published siRNA (red arrows indicate siRNAs with a reported high activity; gray arrows indicate siRNAs with unclear activity; black arrows indicate siRNAs with poor activity). “High activity” was defined as a significant (*p <* 0.05) repression of viral RNA accumulation by ⩾50% in infected cultured cells [53, 55, 57, 59, 61, 62, 58] (and this study), or significant repression of viral plaque count by ⩾50% [54, 60], or a measured IC_50_ for cell infection inhibition being ⩽10 nM [35]. siRNAs have unclear activity when they were assessed in a siRNA mixture rather than individually [56], when they were combined with an anti-hACE2 aptamer before administration to cultured cells [63], or when they were only tested at a 100 nM concentration (gray arrows; also denoted by a question mark for the number of highly active siRNAs in column “High activity siRNAs / total”). SARS-CoV-2 ORF coordinates are from GenBank ACC # OP278726.1; mature proteins generated from the Orf1ab polyprotein are indicated by pink rectangles.

Our siRNA optimization procedure was performed on the January 8, 2021 GISAID variant collection (330,132 sequenced variants, including 311,994 correctly sequenced variants from human donors). In the course of the study, many additional variant sequences (including variants of concern) have been deposited at GISAID: at the time of writing, 13,243,751 variant sequences were available, including 13,033,077 correctly sequenced variants collected from human donors (September 26, 2022 GISAID dataset). Even in this updated variant list, and after excluding unrecognized variants because of sequence ambiguities, our 6 siRNAs with high activity only miss 7,918 to 29,433 variants each, and 12 distinct triplets of siRNAs can recognize each and every one of the *>*13,000,000 sequenced variants (see Figure 5**A**).

**Figure 5.**
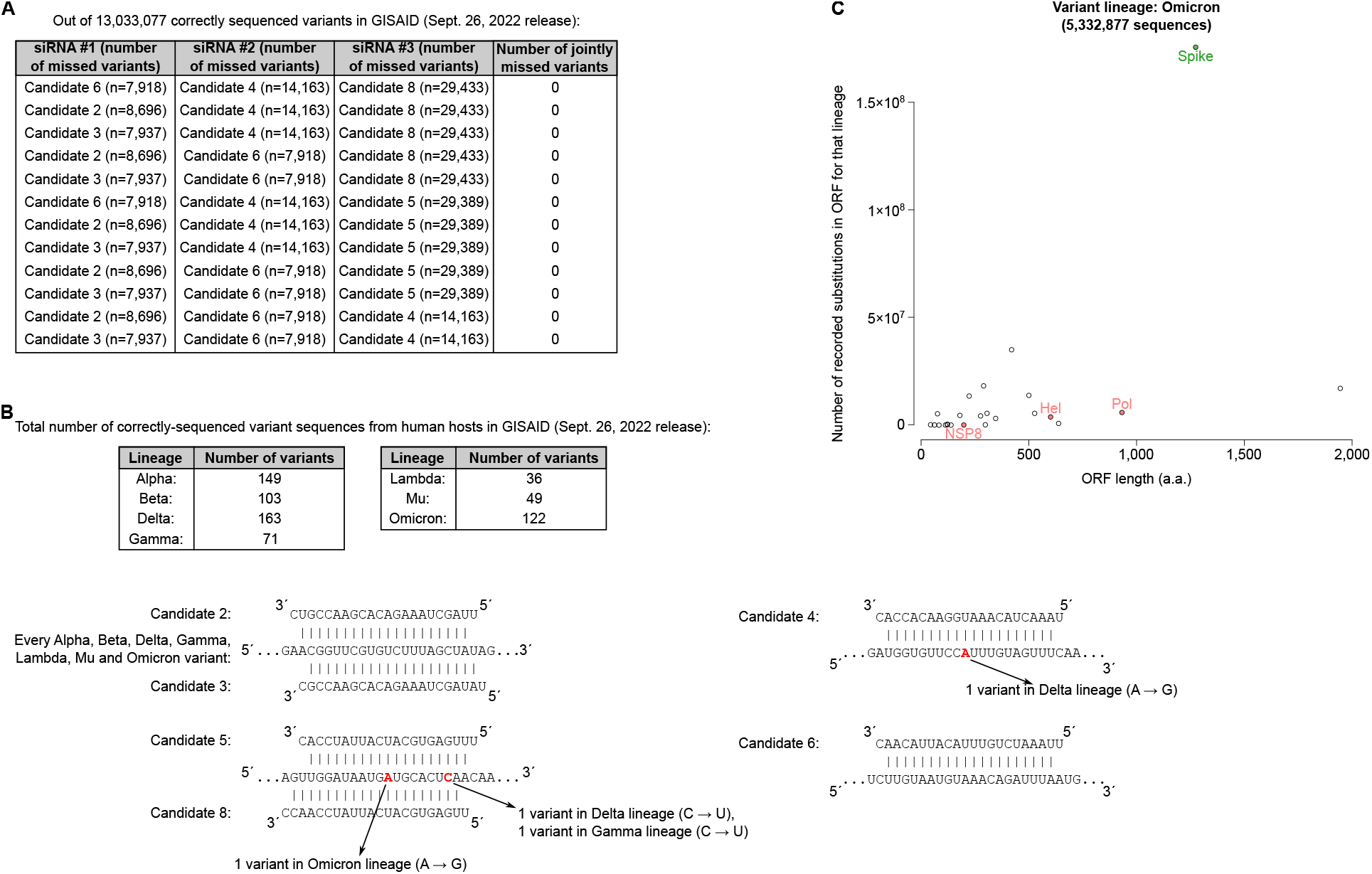
Recognition of newly emerged viral variants. **A**. Among the 20 possible 3-siRNA combinations among our 6 siRNAs with high activity, 12 combinations do not miss any of the correctly sequenced variants (as of September 26, 2022), excluding variants that were not matched perfectly because of sequence ambiguities. A “missed variant” is a variant not exhibiting a perfect 13-mer match to nt 2–14 of the siRNA guide strand. **B**. 7 variant lineages of concern or of interest (Alpha to Omicron) are recorded in the latest GISAID release at the time of writing, with various numbers of deposited sequences (see tables at the top). Each of these variant sequences is perfectly recognized by our siRNA Candidates #2, 3 and 6 and almost all of them are recognized by Candidates #4, 5 and 8 (see predicted RNA duplexes at the bottom). **C**. Number of recorded amino acid substitutions in Omicron lineage variants in GISAID (Sept. 26, 2022 release) for each viral protein (*y* axis) as a function of their length (*x* axis). The Spike protein is highlighted in green, while the three proteins whose gene is targeted by our siRNAs are shown in pink. The total number of Omicron lineage variants in that GISAID release is given above the graph. See Supplementary Figure 4 for other variant lineages of concern or of interest.

Several “variant of concern” lineages have emerged since the beginning of 2021 (*i*.*e*.: SARS-CoV-2 lineages exhibiting higher transmissibility, or more severe symptoms, or substantial escape from previously-acquired immunity, or limited response to existing treatments or diagnostic procedures). Importantly, and despite the *>*18-month-long evolution of the virus in the human population, each of these variants of concern is efficiently recognized by the siRNAs that we designed in January 2021 (see Figure 5**B**). This is essentially due to the fact that such variants tend to accumulate mutations in the *Spike* gene, whose protein product is exposed to the host immune system (hence, under a positive selection pressure) while our siRNAs target genes in Orf1ab (Candidates #2, 3 and 4 target the *Pol* gene, Candidates #5 and 8 target the *Nsp8* gene and Candidate 6 targets the *Hel* gene) (see Figures 4, 5**C** and Supplementary Figure 4).

## DISCUSSION

Despite the prompt development of efficient and safe vaccines against SARS-CoV-2, the COVID-19 outbreak has claimed more than 6.5 million human lives, and caused countless difficulties in the social and economic life. Several treatments have been developed, including anti-inflammatory medication for the most severely affected patients [64], polymerase inhibitors [65] and monoclonal antibodies [66] for nonhospitalized patients.

Besides small molecules and protein-based drugs, the development of siRNAs offers an attractive possibility, provided that suitable administration methods are developed. Biochemical determinants of their repressive activity are very precisely described, allowing efficient design of active siRNAs with limited predicted secondary effects. The great adaptability of siRNA design guarantees a simple and efficient update of siRNA sequences when novel viral variants emerge.

The rapid availability of many viral variant sequences has been a major innovation during the COVID-19 outbreak, and any future viral epidemics is expected to be characterized at least as fast. Using such a variant sequence corpus as the only required input, and applying established rules in siRNA activity, we could design 8 siRNAs with a predicted high antiviral efficiency – with 6 of them indeed proving highly active *ex vivo*. These siRNAs were raised against highly-conserved regions of the SARS-CoV-2 genome as known in January 2021. These regions were not only very constant among the ≈300,000 sequenced variants at that time – they also proved constant since then despite 18 more months of viral circulation in the human population, and our 6 highly active siRNAs still exhibit a perfect 13-mer match to almost every variant as of September 2022 – with 3-siRNA combinations being sufficient to recognize every correctly sequenced variant at that date. It is nonetheless expected that resistant viral variants would probably be selected if such siRNAs were widely administered to the human population. To reduce such risks, it would probably be necessary to track viral variant emergence and to update siRNA design periodically, always administering combinations of siRNAs targeting distinct viral regions (hence: unlikely to mutate in the same variant) rather than a single siRNA.

The main obstacle to siRNA usage *in vivo* lies in the difficulty of delivery to the target organs (in the case of COVID-19: respiratory epithelia). But the administration of a nebulized siRNA solution by inhalation is under development [30] and it gave promising results with chemically-modified or unmodified siRNAs directed against SARS-CoV-2 [61, 58]. Optimal timing for siRNA administration would also need to be optimized (*e*.*g*., administering the siRNAs at various time points before or after viral exposure, then scoring clinical outcomes), because the amplitude of viral replication and its anatonomical localization vary greatly in the course of COVID-19 infection [67]. If a robust method for siRNA administration can be optimized, siRNAs would therefore offer a simple, efficient and easily adaptable therapeutic response in case of future viral pandemics.

## DATA AVAILABILITY STATEMENT

Raw, intermediate and processed data, as well as every script used for the analyses shown in the article, are accessible at https://github.com/HKeyHKey/Houbron_et_al_2023.

## ACKNOWLEDGEMENTS

We gratefully acknowledge all data contributors, *i*.*e*. the Authors and their Originating laboratories responsible for obtaining the specimens, and their Submitting laboratories for generating the genetic sequence and metadata and sharing via the GI-SAID Initiative, on which this research is based. The authors are grateful to Guillaume Canal for his assistance in bioinformatics analyses.

## SUPPLEMENTARY DATA

**Supplementary Figure 1.**
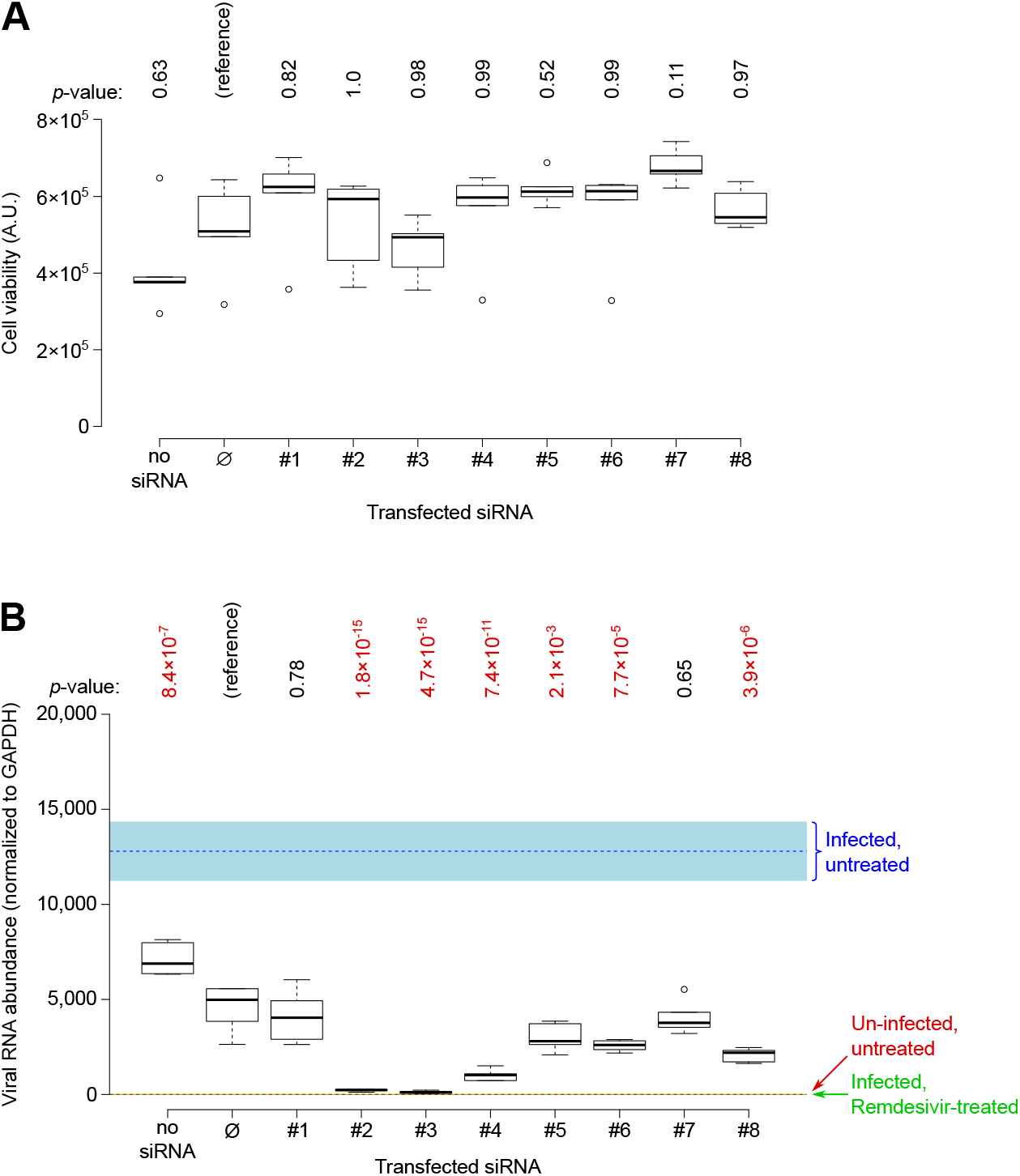
Inocuity and anti-viral efficiency of the 8 candidate siRNAs in A549-hACE2 cells. siRNAs were transfected at 20 nM in A549-hACE2 cells. **A**. Cell viability was assessed 72 h after transfection (6 biological replicates). **B**. Cells were transfected with 20 nM siRNA, infected 7 h later, then viral RNA was quantified by RT-qPCR 48 h after infection (6 biological replicates). Same conventions than in Figure 3. Significance was assessed using Dunnett’s test, with the “∅” siRNA as a reference; *p*-values are indicated above each graph, with *p*-values lower than 0.05 being shown in red.

**Supplementary Figure 2.**
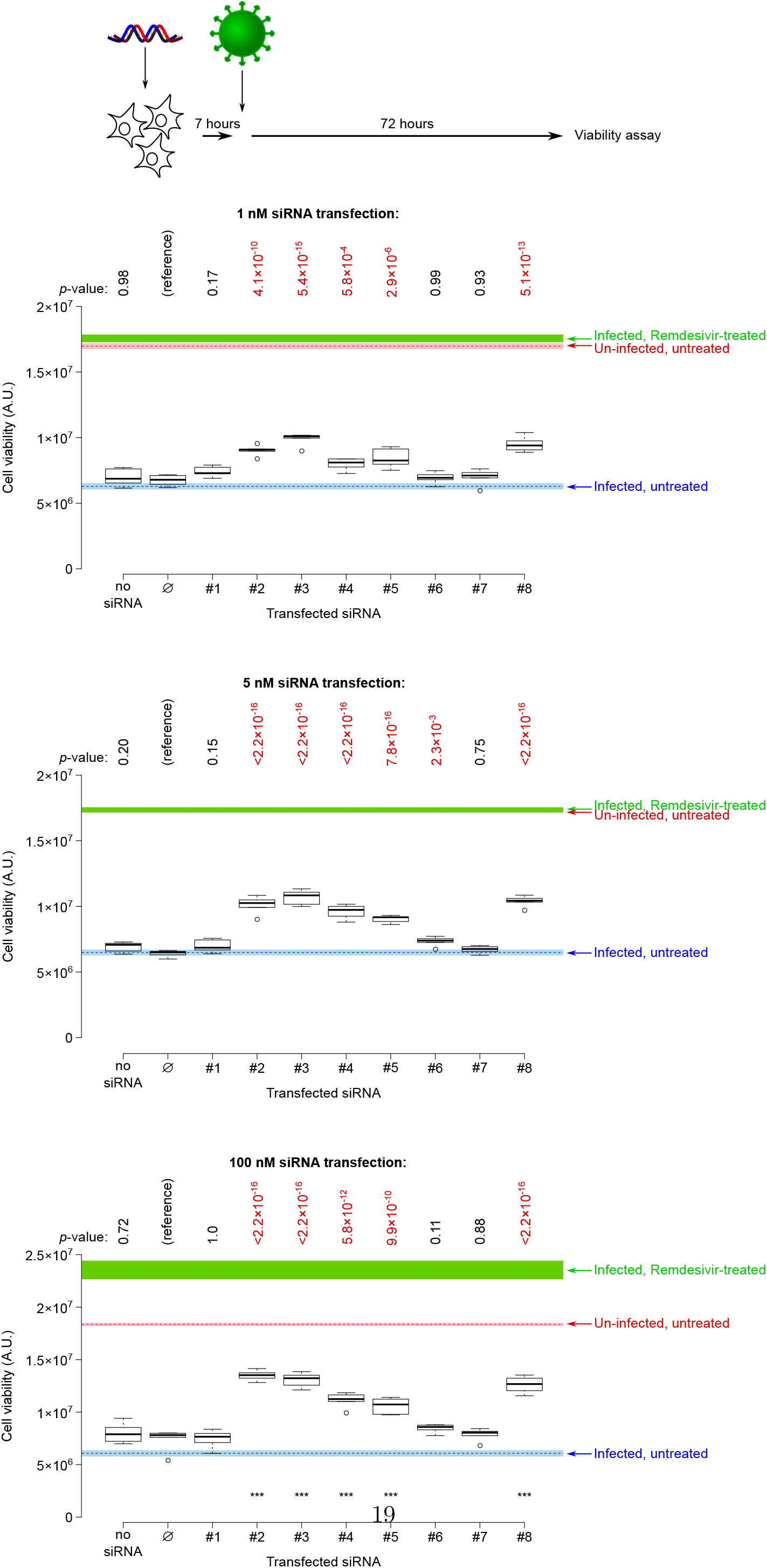
*Ex vivo* assessment of siRNA antiviral activity at various concentrations. Same conventions than in Figure 3**C**.

**Supplementary Figure 3.**
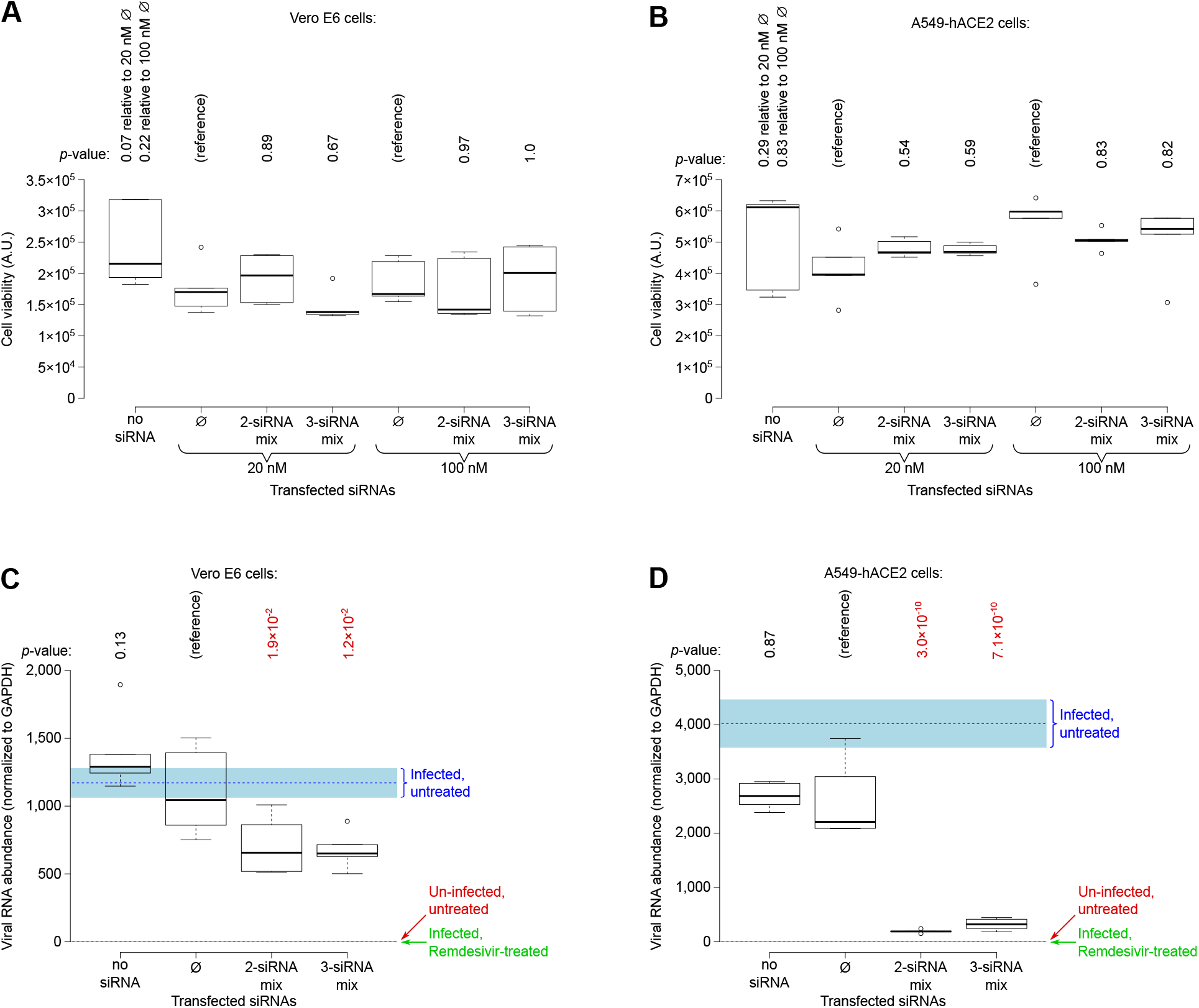
Inocuity and anti-viral efficiency of siRNA combinations in cultured cells. **A, B**. Combinations of 2 siRNAs (siRNAs #3 and 8) or of 3 siRNAs (siRNAs #3, 4 and 8) were transfected at a total concentration of 20 or 100 nM into Vero E6 (panel **A**) or A549-hACE2 (panel **B**) cells, then cell viability was assessed 72 h after transfection (6 biological replicates). **C, D**. siRNA combinations were transfected at 20 nM, infected 7 h later, then viral RNA was quantified by RT-qPCR 48 h after infection (6 biological replicates). Same conventions than in Figure 3. Significance was assessed using Dunnett’s test, with the “∅” siRNA as a reference; *p*-values are indicated above each graph, with *p*-values lower than 0.05 being shown in red.

**Supplementary Figure 4.**
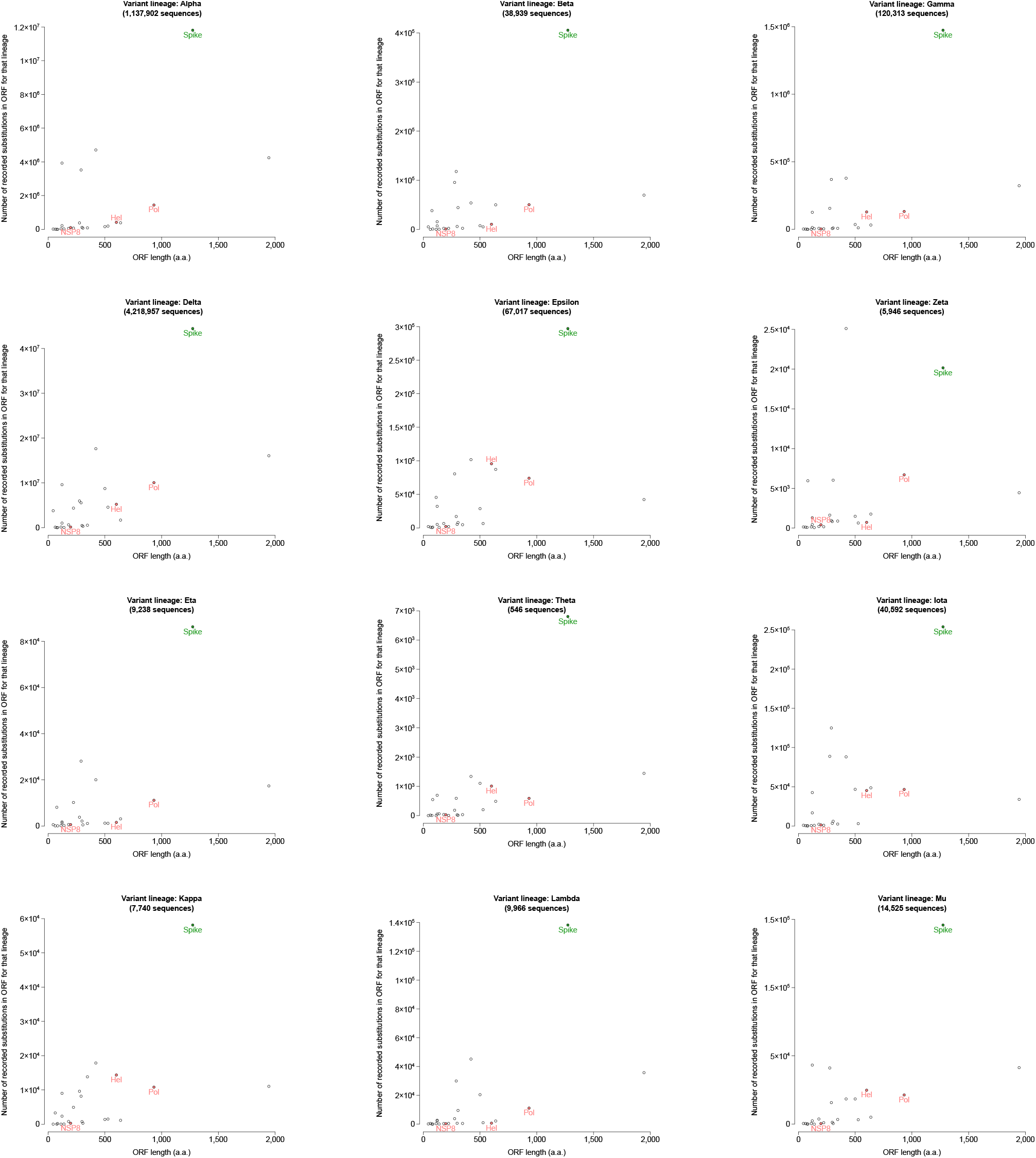
Amino acid substitution per gene in variants of concern and variants of interest. Same conventions than in Figure 5**C**.

